# Fluoxetine effects on behavior and adult hippocampal neurogenesis in female C57BL/6J mice across the estrous cycle

**DOI:** 10.1101/368449

**Authors:** Christine N. Yohn, Sophie Shifman, Alexander Garino, Emma Diethorn, Leshya Bokka, Sandra A. Ashamalla, Benjamin Adam Samuels

**Author notes:** Address correspondence to: Benjamin Adam Samuels; Christine N. Yohn.

## Abstract

Some mood disorders, such as major depressive disorder, are more prevalent in women than in men. However, historically preclinical studies in rodents have a lower inclusion rate of females than males, possibly due to the fact that behavior can be affected by the estrous cycle. Several studies have demonstrated that chronic antidepressant treatment can decrease anxiety-like behaviors and increase adult hippocampal neurogenesis in male rodents. However, very few studies have conclusively looked at the effects of antidepressants on behavior and neurogenesis across the estrous cycle in naturally cycling female rodents. Here we analyze the effects of chronic treatment with the selective serotonin reuptake inhibitor (SSRI) fluoxetine (Prozac) on behavior and adult hippocampal neurogenesis in naturally cycling C57BL/6J females across all four phases of the estrous cycle. Interestingly, we find that the effects of fluoxetine on both behavior and adult hippocampal neurogenesis are driven by mice specifically in the estrus or diestrus phases of the estrous cycle. Taken together our data is the first to illustrate the impact of fluoxetine on brain and behavior across all four stages of the murine estrous cycle.

**Highlights:**

- Chronic fluoxetine reduces anxiety-like behaviors in naturally cycling female mice
- Chronic fluoxetine increases adult hippocampal neurogenesis in naturally cycling female mice
- The effects of chronic fluoxetine on behavior and adult hippocampal neurogenesis are driven by the estrus and diestrus phases of the estrous cycle

## 1. Introduction

Although major depressive disorder is more prevalent in women than men (Kessler, 2003; Sloan & Kornstein, 2003), females are often excluded from rodent experimental studies since fluctuations in ovarian steroid hormones (Arakawa et al., 2014; Lovick, 2012), such as estrogen, estradiol, and progesterone, across the female’s estrous cycle can confound experimental results. In humans, women can experience depression and anxiety due to premenstrual syndrome, where variations in mood states are correlated with different secretion patterns of estrogen and progesterone across the menstrual cycle (Shors & Leuner, 2003). Rodents display similar fluctuations in behavior, with diestrus female rodents having higher responses to stress and increased anxiety-related behaviors as compared to proestrus females (Lovick, 2012; D’Souza & Sadanada, 2017; Sayin et al., 2014; Marcondes et al., 2001). While these studies illustrate the influence of the estrous cycle on animal behavior, they were predominately conducted in rats not mice. In comparing two mouse strains, Meziane and colleagues (2007) observed that C57BL/6J females have less variation in anxiety-like behaviors across the estrous cycle than BALB/cByJ females. However, commonly used negative valence tests associated with anxiety-like behavior, such as elevated plus maze (EPM) and novelty suppressed feeding (NSF), were not included in this study. Therefore, detailed behavioral analyses across the mouse estrous cycle is still needed.

Variations in gonadal steroid hormone secretion patterns, as seen in cycling females, can contribute to hippocampal structural and functional impairments, such as alterations in adult neurogenesis (Tanapat et al., 2005; Barha et al., 2009) within the subgranular zone of the dentate gyrus (DG) (Tanapat et al., 1999). For instance, proestrus rats have higher DG cell proliferation than estrus or diestrus rats with these differences most likely attributed to higher estradiol levels during the proestrus phase (Sadrollahi et al., 2014; Tanapat et al., 1999; Pawluski et al., 2009). While many studies document the effects of the estrous cycle and administration of estrogens to mimic a cycle stage in ovariectomized (OVX) rats on DG cell proliferation, these effects may be species specific since Lagace and colleagues (2007) observed no significant differences in adult hippocampal cell proliferation within C57BL/6J mice. However, whether the estrous cycle impacts other stages of adult hippocampal neurogenesis, such as doublecortin which labels young and mature neurons (Plumpe et al., 2006), in C57BL/6J mice is unknown. Regardless of species differences in estrous effects on adult hippocampal cell proliferation, understanding the impact the estrous cycle has on adult hippocampal neurogenesis is important since pharmacotherapies, such as antidepressants, exert beneficial effects on behavior in part by increasing adult hippocampal neurogenesis (Malberg et al., 2000; David et al., 2009; Santarelli et al., 2003). While Sayin and colleagues (2014) illustrate that citalopram, a selective serotonin reuptake inhibitor (SSRI), alleviates differences in anxiety-like behavior between proestrus and non-proestrus rats, few studies have examined the impact of SSRIs on behavior and neurogenesis in intact, cycling female mice. Therefore, the current study aims to assess the behavioral and neural effects of fluoxetine (FLX), a SSRI, across the estrous cycle.

## 2. Methods

### 2.1 Subjects

Adult 8 week old female C57BL/6J strain mice (n = 41) were purchased from Jackson laboratories. All mice were maintained on a 12L:12D schedule with food and water provided *ad libitum*. For 3 weeks FLX (18 mg/kg/day) or vehicle (deionized water) was delivered via oral gavage. On behavioral testing days FLX or vehicle was administrated after mice completed the behavioral test to avoid acute effects. All testing was conducted in compliance with the NIH laboratory animal care guidelines and approved by Rutgers University Institutional Animal Care and Use Committee.

### 2.2 Vaginal Lavage

To examine estrous cycle state vaginal lavages were performed throughout FLX/vehicle treatment, after completing each behavioral test, and prior euthanasia. In order to collect the samples, a pipet was filled with ddH_2_O, placed at the opening of the mouse’s vaginal canal (without penetration) with ddH_2_O gently expelled and suctioned back (for detailed methods see McLean et al., 2012; Byers et al., 2012). Samples were then placed on a slide warmer to dry for approximately 5 minutes and imaged under a EVOS FL Auto 2.0 microscope (Thermofisher Scientific) at 10x magnification. Estrous phase was identified by the presence or absence of nucleated epithelial cells, cornified epithelial cells, and leukocytes (Byers et al., 2012; Felicio et al., 1984). Mice in proestrus displayed mostly nucleated and some cornified cells (Figure 1B). Estrus was recorded as the presence of mostly cornified epithelial cells, with the presence of a few nucleated cells in early estrus (Figure 1B). Metestrus was determined by the presence of cornified epithelial cells and polymorphonuclear leukocytes (Byers et al., 2012), while mice in diestrus contained mainly polymorphonuclear leukocytes with few epithelial cells being present (Figure 1B).

**Figure 1.**
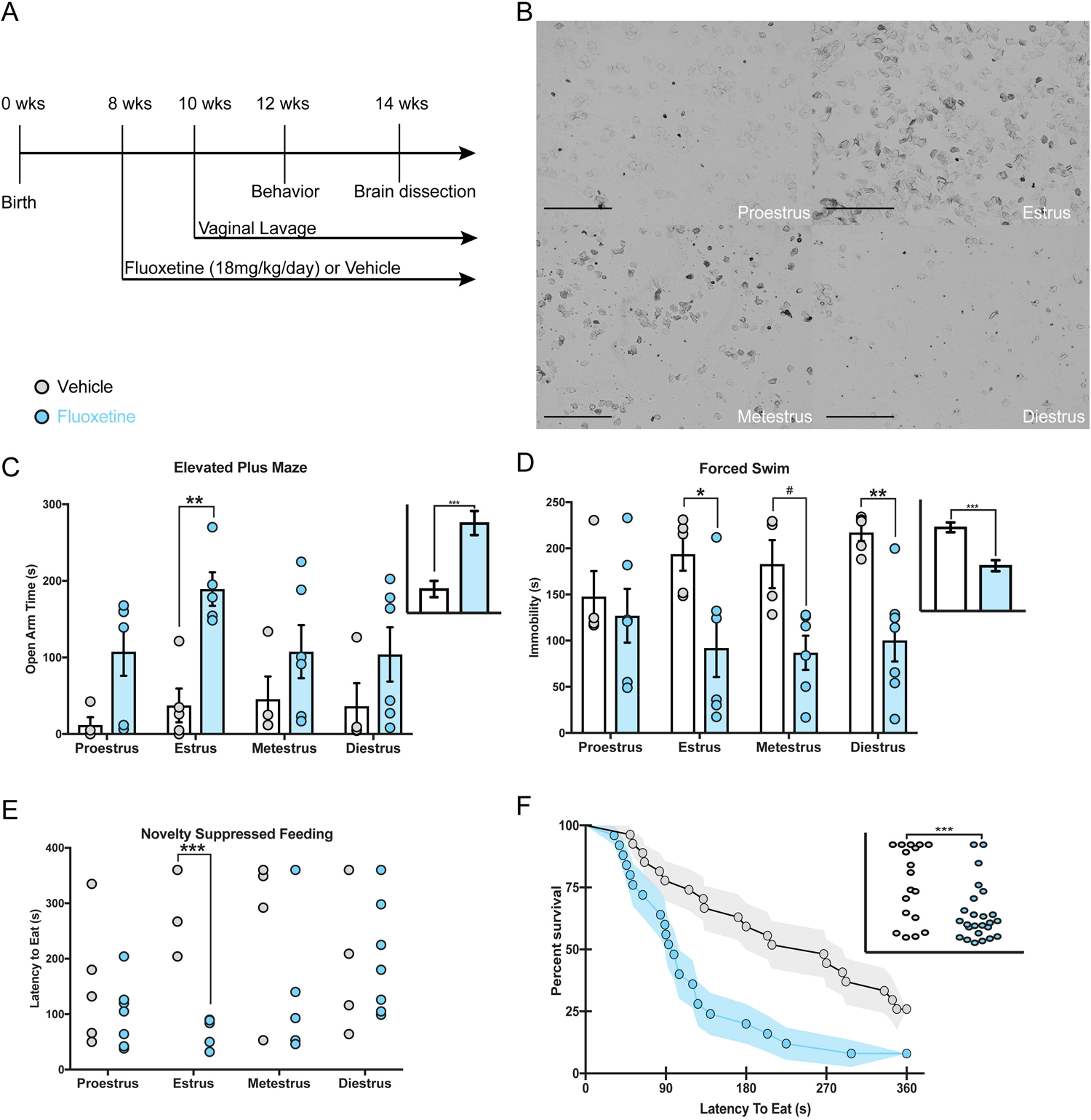
Behavioral differences between FLX- and vehicle-treated female mice are also mediated by estrous phase. A timeline depicting the experimental study is depicted above (A) with representative images of the four stages of the estrous cycle (10x magnification; scale = 500 µm; B). Separate 2×4 ANOVAs were run and revealed that overall females treated with FLX spend less time in the open arms than vehicle mice (p < 0.001), with this group difference being evident within the estrus phase of the estrous cycle (p = 0.005; C). In the FST, FLX appeared to reduce overall immobility time (p < 0.001), with FLX-treated females in the estrus and diestrus phases having significantly lower immobility scores than vehicle-treated females (D). Also, FLX-treated females have quicker latencies to eat in the NSF task than vehicle-treated females, with group differences being evident within the estrus phase (B). * denotes p < 0.05; ** denotes p <0.01; *** denotes p <0.001

### 2.3 Behavioral Testing

#### 2.3.1 Open Field (OF)

Motor activity was quantified in five Plexiglass open field boxes 43 × 43 cm^2^ (Kinder Scientific). The recording of x-y ambulatory movements was recorded by two sets of 16 pulse-modulated infrared photobeams placed on opposite walls 2.5 cm apart to. As previously described (David et al., 2009), activity chambers were computer interfaced for data sampling at 100ms resolution. The computer software predefines grid lines that divide each open field chamber into center and periphery regions, with the center being a square 11cm from the wall. The number of entries, distance traveled, and total time spent in the center were recorded, as well as percent of distance traveled in the center defined as center distance divided by total distance traveled (Supplemental Figure 1A). To measure overall motor activity total distance (cm) was quantified.

#### 2.3.2 Novelty Suppressed Feeding (NSF)

After undergoing 18 hours of food deprivation within their home cage, mice were placed in the corner of a testing apparatus (50×50×20 cm) filled with approximately 2 cm of corncob bedding and a single pellet of food attached to a white platform in the center of the box. The center of the box was illuminated at 1500 lux. The NSF test lasted 6 minutes, where the latency to eat (defined as the mouse sitting on its haunches and biting the pellet with the use of forepaws) was timed/recorded. If a mouse did not consume food during the NSF a latency of 360secs was recorded. Immediately afterwards, the mice were transferred to their home cage to assess home cage feeding behavior for 5 minutes (Supplemental Figure 1D-F). During this task latency to eat and amount of food consumed was measured as a control for feeding behavior observed in the NSF task. Each mouse was weighed before food deprivation and after home cage feeding to assess the percentage of body weight loss. Following home cage feeding, mice were placed in a new home cage with cage mates and returned to the colony room.

#### 2.3.3 Light Dark (LD)

The light/dark test was conducted in an open field chamber measuring 43.2 × 43.2 cm (Kinder Scientific, USA), with a clear floor and walls. To divide the open field into separate light and dark compartments, a dark plastic box that covered one third of the chamber was inserted. The dark box was opaque to visible light, but transparent to infrared light, and contained an opening that allowed passage between the light and dark compartments (David et al., 2009). The light compartment was brightly illuminated at 1000 lux. At the beginning of each 5-minute test, mice were placed in the dark compartment. An observer blind to treatment groups recorded the latency to emerge into the light (Supplemental Figure 1C). Using Activity Monitor (Kinder Scientific, USA) software, total time in the light and ambulatory distance in both compartments was analyzed. To calculate percent distance traveled in the light, distance traveled in the light was divided by total distance traveled (Supplemental Figure 1B).

#### 2.3.4 Elevated Plus Maze (EPM)

The EPM test consisted of a plus-shaped apparatus with two open and two closed arms (side walls), elevated 2 feet from the floor. During the five-minute test, the mouse’s behavior was recorded from a video camera mounted above each EPM arena. EthoVision (Noldus) software was then used to score time spent in the open arms, entries, and distance traveled in both open and closed arms. By dividing total open arm distance traveled by total distance traveled, we were able to analyze percent distance traveled in open arms.

#### 2.3.5 Forced Swim Test (FST)

A modified FST procedure suitable for mice was used (David et al., 2009). Individual cylinders (46 cm tall × 32 cm in diameter × 30 cm deep) were filled with room-temperature water (25–26°C) before placing each mouse into the cylinder. Two sets of photobeams were mounted on opposite sides of the cylinder (Kinder Scientific, USA) to allow for the recording of swimming behavior during the 6-minute test. Immobility times (measured by beam breaks over 5-second intervals) were assessed during the last 4-minutes of the test since mice are habituating to the task during the initial 2-minutes of the test.

### 2.4 Brain Collection, Sectioning and Immunohistochemistry

#### 2.4.1 Brain Collection and Sectioning

Subsequent to the completion of all behavioral testing, brains were collected from all experimental mice. Mice were anesthetized with ketamine (80mg/kg) and perfused transcardially with PBS followed by 4% paraformaldehyde. Brains were collected and stored in 4% paraformaldehyde overnight at 4°C. Next, brains were switched to a 30% sucrose 0.1% sodium azide (NaN_3_) in PBS solution and stored at 4°C until they were sectioned. Using a cryostat, serial sections of the hippocampus (Franklin & Paxinos mouse brain atlas 3^rd^ edition; Bregma –1.22 to –3.88) were collected onto Superfrost Plus slides (Thermofisher Scientific) and stored at –20°C until staining and further analysis.

#### 2.4.2 Ki67 labeling for cell proliferation

The effects of FLX treatment on cell proliferation were assessed across 12 serial sections of the hippocampus. Using mailers, slides were washed in 1% Triton X-100 PBS for 5 minutes before undergoing three PBS washes. Next, slides were incubated in warm citrate buffer for 30 minutes and then washed times in PBS. Slides were then transferred to an opaque moisture chamber (details) for the blocking and overnight incubation step. Slides were blocked for 1 hour in 10% normal goat serum (NGS) diluted in PBS before being incubated overnight at 4°C in anti-rabbit ki67 (1:500; abCam, ab16667) diluted in 2% NGS PBS. Following 18 hours of incubation slides were washed 3 times in PBS before being incubated at room temperature for 2 hours in CY-5 goat anti-rabbit (1:1000, Invitrogen, Thermo Fisher Scientific, A10523) diluted in 2% NGS PBS. Next slides were washed with PBS then counterstained with DAPI (1:15000) for 15 minutes. Finally, slides were washed with PBS and cover slipped using the mounting medium prolong diamond. Fluorescent images were taken using a EVOS FL Auto 2.0 microscope (Thermofisher Scientific) at 10x magnification, where ki67^+^ cells overlayed with DAPI across the 12 sections of hippocampus were collected and counted.

#### 2.4.3 Doublecortin (DCX) labeling for maturation

12 serial hippocampal sections for doublecortin (DCX) were stained using the primary antibody doublecortin anti-goat (1:500; Life technologies; 481200) and secondary antibody CY-5 goat anti-rabbit (1:1000, Invitrogen, Thermo Fisher Scientific, A10523). Fluorescent images were taken using using a EVOS FL Auto 2.0 microscope (Thermofisher Scientific) at 10x magnification, where DCX^+^ cells across the 12 sections of hippocampus were collected and counted. Following imaging, DCX^+^ cells were counted and subcategorized according to their dendritic morphology: DCX^+^ cells with no tertiary dendritic processes and DCX^+^ cells with complex, tertiary dendrites. The maturation index was defined as the ratio of DCX^+^ cells possessing tertiary dendrites over the total DCX^+^ cells.

### 2.5 Statistical Analyses

To analyze both behavioral and molecular differences between treatment groups and estrous cycle stages separate 2×4 analyses of variance (ANOVA) were conducted. Each ANOVA was followed by subsequent Bonferroni post-hoc analyses to further assess between and within group differences across the estrous cycle. Since we imposed a cutoff time during the NSF, we ran a Kaplan Meier survival analysis (nonparametric test) that permits censoring of these data points to analyze differences in feeding latencies.

## 3. Results

### 3.1 FLX and vehicle behavioral differences across estrous cycle

We treated a large cohort of adult C57BL/6J female mice (n= 41) with 18mg/kg FLX for three weeks (Figure 1A) and then exposed these mice to the Open Field (OF), Light Dark Test (LD), EPM, NSF test, and Forced Swim Test (FST). To assess estrous cycle phase (Figure 1B), vaginal lavages were performed two weeks prior to behavior, following each behavioral test, and on the days between behavioral tests. We found significant treatment effects in the EPM and FST, such that FLX mice spent more time in the open arms (F(1,32)=19.05, p <0.001; Figure 1C) and less time immobile (F(1,35) = 15.15, p < 0.001; Figure 1D) than vehicle mice. In the EPM, planned Bonferroni post-hoc comparisons revealed treatment differences in open arm time within the estrus phase, such that estrus FLX-treated mice spent more time in the open arm than estrus vehicle-treated mice, t(32) = 1.56, p = 0.001 (Figure 1C). Additionally, post-hoc comparisons revealed immobility time in the FST was significantly different between treatment groups within the estrus and diestrus phases, such that vehicle mice spent significantly more time immobile than FLX mice within both the estrus (t(10) = 2.41, p = 0.02) and diestrus (t(11) = 2.23, p = 0.03) phases (Figure 1D), with a nonsignificant trend emerging in the metestrus phase (t(9) = 2.62, p = 0.052). We found no significant effects of treatment or estrous phase nor interaction effect on behaviors within the OF and LD tests (Supplemental Figure 1A-C).

In the NSF, a Kaplan Meier survival analysis log rank test revealed that FLX significantly reduces latency to feed, x^2^(1) = 8.37, p = 0.0038 (Figure 1E, 1F), as compared to vehicle mice. Additionally, we observed that the estrus phase of the estrous cycle impacted group differences in latency to feed, with FLX females in estrus having lower latencies to feed than estrus vehicle females x^2^(1) = 6.9, p = 0.008. There was no treatment effect nor estrous effect on home cage latency, amount of food consumed in home cage, or percent weight change (Supplemental Figure 1D-F).

Taken together, these data suggest that the effects of FLX on behavior are most consistent across tests during the estrus stage.

### 3.2 Fluoxetine treatment and estrous cycle state impact adult neurogenesis

Several days following the FST, we perfused the mice and collected serial sections through the DG. Next, we performed immunostaining to determine the effects of FLX on the distinct stages of adult hippocampal neurogenesis across the different phases of the estrous cycle. We first stained for the cell proliferation marker Ki67 and found that FLX-treated mice had more Ki67^+^ cells than vehicle-treated mice F(1, 37) = 5.34, p = 0.026 (Figure 2A). We observed no estrous cycle effects nor interaction on number of Ki67^+^ cells (p’s > 0.05).

**Figure 2.**
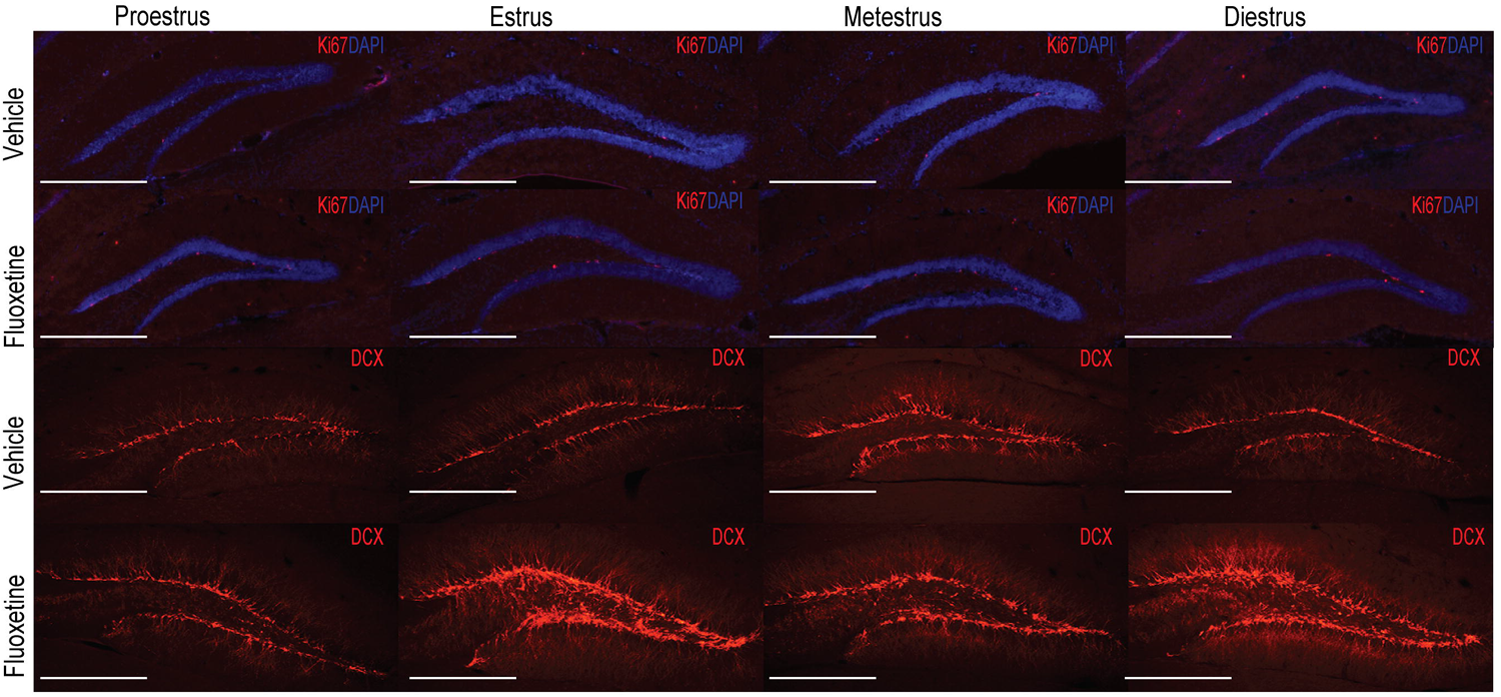
FLX increases all stages of neurogenesis with differences most pronounced within both the estrus and diestrus phases. Visual representation of Ki67 and DCX immunostaining is depicted above for each treatment group across the estrous cycle phases. Images were taken at 10x magnification (scale = 500 µm). FLX-treated females had higher adult cell proliferation (Ki67^+^) compared to vehicle animals, with no differences between groups emerging across the estrous cycle (A). FLX-treated females had more immature (B) and mature (C) neurons as well as a higher maturation index (D) compared to vehicle-treated females, with differences between groups most pronounced during the estrus and diestrus phases. * denotes p < 0.05; ** denotes p <0.01; *** denotes p <0.001

We next performed immunostaining with the young neuron marker DCX, and observed that FLX-treated mice showed more DCX^+^ cells within the DG than vehicle-treated mice F(1, 37) = 13.44, p < 0.001. We found a significant interaction effect between treatment and estrous phase F(3, 37) = 4.57, p = 0.008. Planned post-hoc comparisons revealed that FLX-treated mice had more DCX^+^ cells than vehicle mice within the estrus (t(9) = 3.95, p < 0.001) and diestrus (t(11) = 3.29, p = 0.009; Figure 2B) phases. Within the vehicle group, planned comparisons revealed that proestrus females had significantly more DCX^+^ cells than both estrus (t(10) = 4.76, p = 0.001) and diestrus (t(11) = 3.71, p = 0.01) female mice (Figure 3A). Additionally, metestrus vehicle-treated mice had significantly more DCX^+^ cells than both estrus (t(9) = 4.8, p < 0.001) and diestrus (t(9) = 3.82, p = 0.008) vehicle-treated mice (Figure 3A). Within the FLX group no differences in DCX^+^ cell expression was observed across the estrous cycle.

To assess maturation of the young neurons, we counted the DCX+ neurons that displayed tertiary dendrites. As expected, FLX-treated mice had more mature neurons than vehicle mice as indicated by the number of DCX^+^ cells with tertiary dendrites (F(1, 37) = 20.56, p <0.001; Figure 2C). A significant interaction between treatment group and estrous cycle phase emerged (F(3, 37) = 5.33, p = 0.004), with FLX mice having more DCX^+^ cells with tertiary dendrites than vehicle mice in the estrus (t(9) = 4.19, p < 0.001) and diestrus (t(11) = 4.23, p < 0.001) phases. Within the vehicle group, planned post-hoc comparisons revealed that proestrus females have more DCX^+^ cells with tertiary dendrites than estrus (t(10) = 6.01, p = 0.003) and diestrus (t(10) = 5.09, p = 0.011) females (Figure 3B). Planned comparisons also revealed that metestrous vehicle-treated females had more mature neurons than estrus (t(9) = 4.94, p = 0.013) and diestrus (t(9) = 4.06, p = 0.04) vehicle-treated females (Figure 3B). Within the FLX group, we observed no differences in expression of DCX^+^ cell with tertiary dendrites across the estrous cycle.

**Figure 3.**
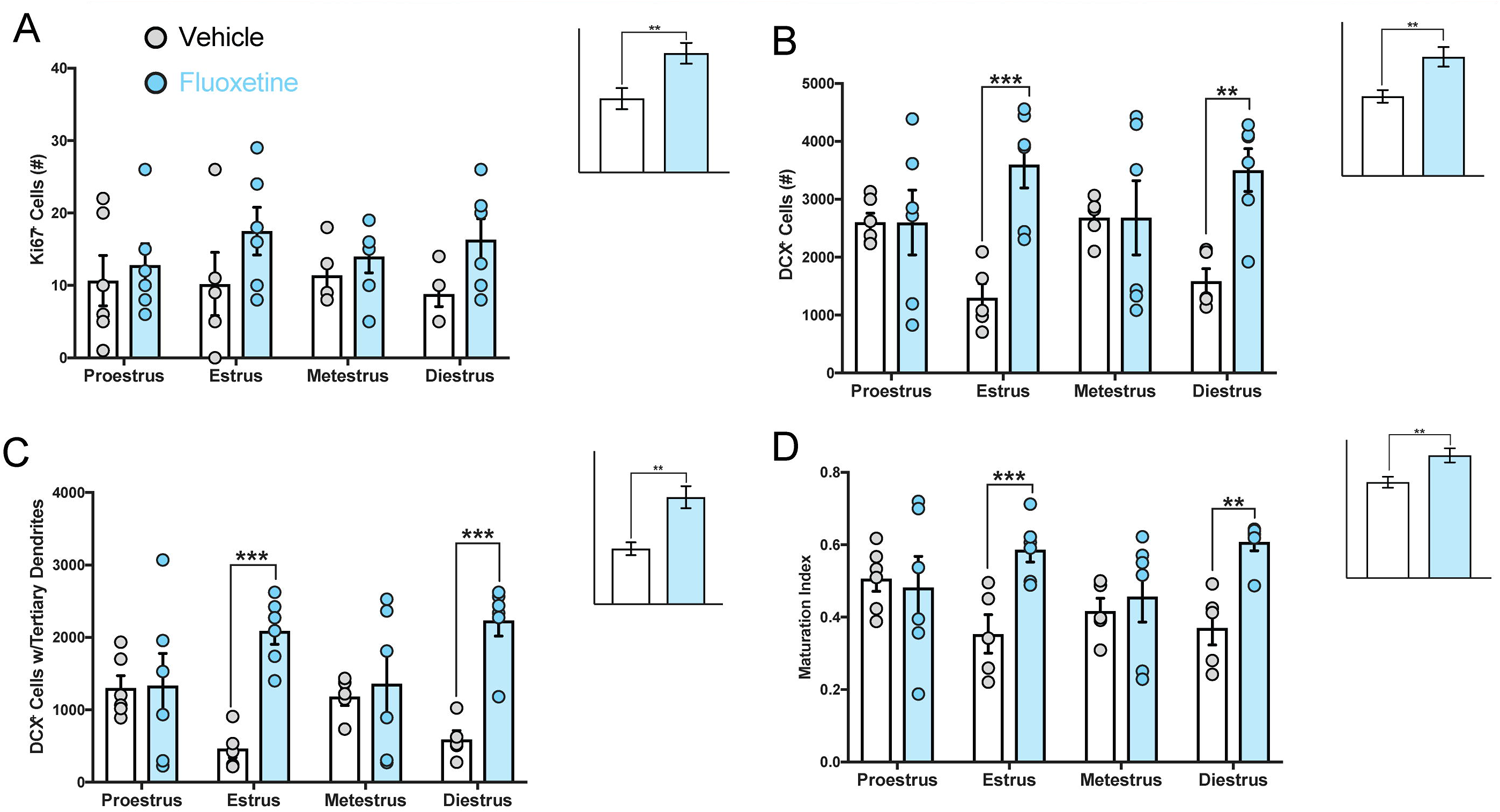
Estrous cycle impacts adult hippocampal neurogenesis in intact, cycling vehicle-treated female mice. Planned post-hoc analyses within the vehicle group was conducted to assess impact of estrous cycle on adult hippocampal neurogenesis. No differences in adult hippocampal cell proliferation was noted across the estrous cycle in vehicle females (A). Proestrus and metestrus females had higher expression of immature (B) and mature (C) neurons than estrus and diestrus females. There was no impact of estrous cycle on maturation index (D). * denotes p < 0.05; ** denotes p <0.01; *** denotes p <0.001

Lastly, we observed that FLX mice have a higher maturation index (F(1, 37) = 10.98, p = 0.002; Figure 2D) than vehicle mice. We observed a significant interaction effect between treatment group and estrous cycle phase (F(3, 37) = 3.23, p = 0.033), with FLX females having a significantly higher maturation index than vehicle females in both the estrus, (t(9) = 3.07, p = 0.004) and diestrus (t(11) = 3.13, p = 0.003) phases (Figure 2D). Within both the vehicle (Figure 3C) and FLX group we observed no differences in the maturation index across the estrous cycle. Taken together, these data suggest that the effects of fluoxetine on adult hippocampal neurogenesis are most pronounced during both the estrus and diestrus stage.

## 4. Discussion

### 4.1 FLX and vehicle behavioral differences across estrous cycle

We used an array of negative valence behavioral tests to evaluate the impact of antidepressant treatment on anxiety-like behaviors across the estrous cycle. Prior to behavioral testing we tracked the females estrous cycle for two weeks to assess whether females were cycling together within the same housing room. Although Meziane and colleagues (2007) observed that females within the same room cycle together, we observed that females within this study had out of sync cycles allowing for us to investigate the different estrous cycle phases during each behavioral test. Similar to previous studies in males (David et al., 2009), our data illustrates that FLX treatment in females reduces anxiety-like behaviors within the EPM and NSF, and decreases immobility in the FST. However, our data demonstrates that estrous cycle significantly impacts the effects of FLX on behavior. FLX-treated females in the estrus phase display a reduction in anxiety-like behavior in the EPM and NSF, and reduced immobility in the FST relative to vehicle-treated estrus females. Diestrus FLX-treated females had lower immobility times in the FST than diestrus vehicle females, but FLX was ineffective in the anxiety-related EPM and NSF tasks. Furthermore, the behavioral effects of FLX were not significant in any of these tasks during metestrus and proestrus. Recently, Sayin and colleagues (2014) observed that proestrus rats display less anxiety-like behaviors than non-proestrus rats in the EPM, with estrous cycle differences attenuated following citalopram administration. Compared to our data, these data suggest that species differences may exist in anxiety-like behaviors across the estrous cycle. Although we did not observe an effect of the estrous cycle on behavior within treatment, a previous study observed that C57BL/6J females have less variation in anxiety-like behaviors across the estrous cycle than BALB/cByJ females (Meziane et al., 2007). Despite this, our data illustrate that FLX treatment has significant effects on behavior relative to vehicle treatment. However, our detailed analyses suggest that these effects of FLX are mainly driven by females in the estrus and diestrus phase.

Differences in behavior between treatment groups within estrus and diestrus may be related to fluctuations in estradiol and progesterone levels (Pawluski et al., 2009; Lovick, 2012). Specifically, estradiol levels are the lowest during estrus and diestrus (Pawluski et al., 2009; Wood et al., 2007). Exogenous estradiol treatment to mimic diestrus in OVX rats results in decreases in anxiety-like behavior in the EPM compared to non-estrogen treated freely cycling diestrus rats (Marcondes et al., 2001). In mice, females in the estrus and diestrus phases are more susceptible to individual housing stress and spend less time in the center of the open field arena than proestrus mice (Palanza et al., 2001). The impact of FLX administration on ovarian steroid hormones across the estrous cycle is understudied in rodents, and future studies will need to assess the relationship between antidepressant and endogenous ovarian hormone levels.

### 4.2 Fluoxetine treatment and estrous cycle state impact adult neurogenesis

Similar to several other studies (Pawluski et al., 2014; Lagace et al., 2007), we observed that chronic FLX administration increased adult hippocampal neurogenesis levels relative to vehicle-treated females. Females treated with FLX had higher levels of cell proliferation (Figure 2A), higher numbers of both immature (Figure 2B) and mature neurons (Figure 2C), as well as a higher maturation index (Figure 2D) than vehicle treated females. However, similar to the effects on behavior, our study is the first to illustrate that the effects of FLX on adult hippocampal neurogenesis are most pronounced during the estrus and diestrus phases. Differences in adult hippocampal neurogenesis within estrus and diestrus could be attributed to natural low-levels of estradiol in these phases. Estrogens, such as estradiol, impact both cell proliferation and cell survival in the DG (Ormerod et al., 2003; Barha et al., 2009) and more proliferating cells are found in the proestrus phase than the non-proestrus phases (Sadrollahi et al., 2014; Tanapat et al., 1999; Pawluski et al., 2009). Discrepancies in proliferating cell numbers across the estrous cycle can be attributed to endogenous estrogen levels naturally peaking during the proestrus phase and decreasing during the estrus and diestrus phase (Pawluski et al., 2009). However, Lagace and colleagues (2007) show that endogenous levels of estradiol do not appear to impact adult hippocampal cell proliferation in mice, since OVX female mice have similar number of proliferating cells (BrdU^+^) and immature neurons (DCX^+^) in the hippocampus as intact female mice. Furthermore, in assessing cell proliferation across 3 phases of the estrous cycle (proestrus, estrus, diestrus), Lagace and colleagues (2007) observed no differences in cell proliferation in the different phases. Similar to Lagace and colleagues (2007), we show that estrous cycle phase does not impact DG cell proliferation levels within treatment group. However, Lagace and colleagues (2007) did not assess differences in immature and mature neurons across the estrous cycle. We found that proestrus and metestrus vehicle-treated female mice have higher numbers of immature neurons (DCX^+^) within the DG than estrus and diestrus vehicle-treated female mice (Figure 3A). Additionally, we show that estrus and diestrus vehicle treated female mice have fewer mature neurons in the DG than proestrus and metestrus vehicle-treated female mice (Figure 3B). Interestingly, FLX specifically increased the numbers of immature (Figure 2B) and mature (Figure 2C) neurons in the DG during estrus and diestrus relative to vehicle-treated mice. Therefore, no differences across the phases of the estrous cycle were observed within the FLX group. These data suggest that the effects of FLX on adult hippocampal neurogenesis are mainly driven by females in the estrus and diestrus phases of the estrous cycle.

Overall our study illustrates that the tracking the estrous cycle in experimental studies is crucial since different estrous phases show significant differences in behavior and adult hippocampal neurogenesis. Furthermore, the effects of FLX treatment are mainly driven by females in the estrus and diestrus phases. Future studies should assess whether antidepressants influence endogenous levels of ovarian hormones, such as estrogen and progesterone, since fluctuations in these hormones across the estrous cycle may attribute to differences in behavior and adult hippocampal neurogenesis. Given that sex differences in the etiologies of mood disorders and symptomologies exist, preclinical studies that determine differences across the estrous cycle are critical for developing a better understanding of how these disorders develop and should be treated in females.

## Acknowledgments

This work was supported by the National Institute of Mental Health [Grant Number R01 MH112861 to BAS]. The authors would also like to thank Mark M. Gergues, Kylee Shivok, and Debbie Ma for their assistance with this project.

---

**Supplemental Figure 1.**
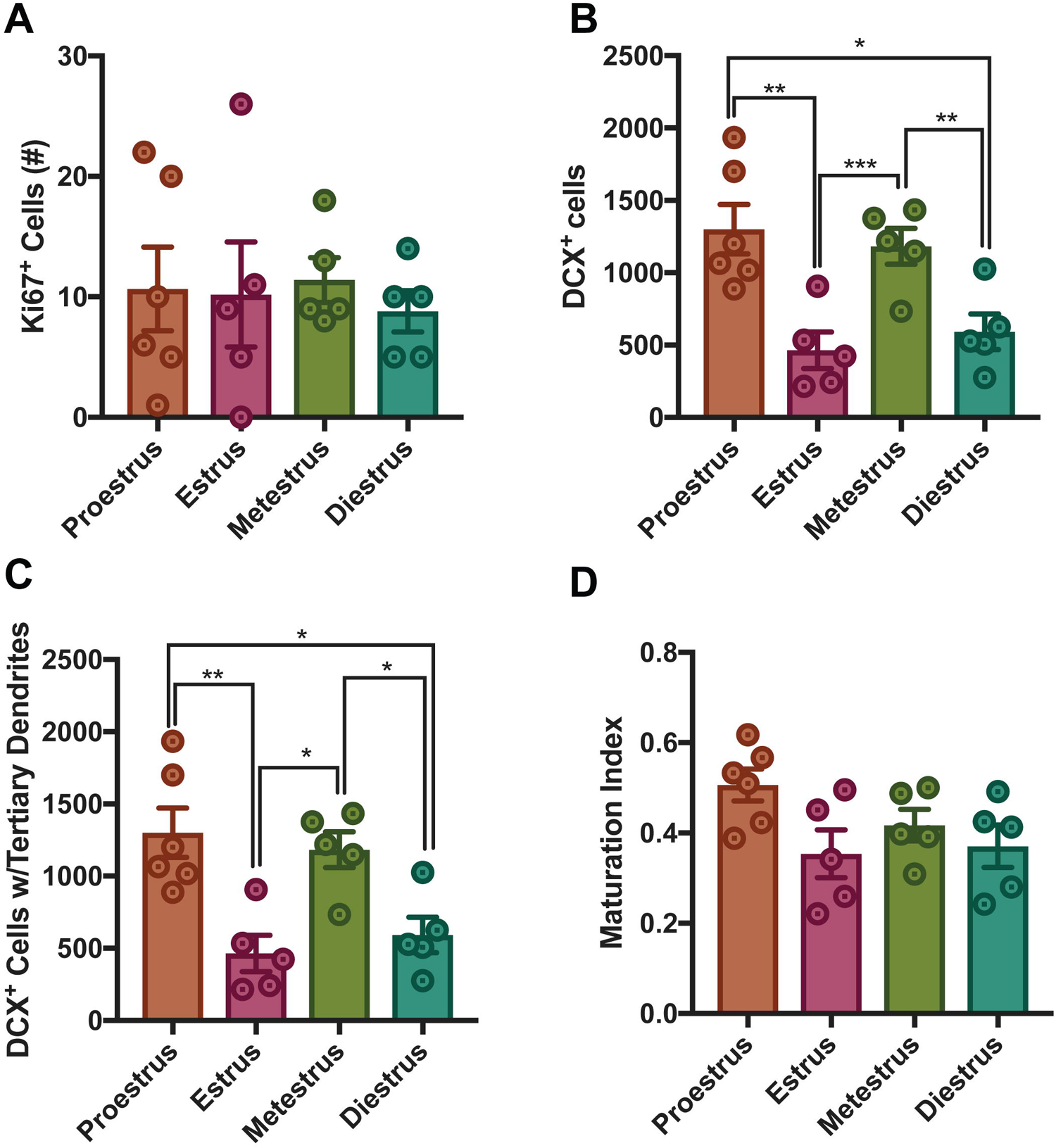
Treatment and estrous cycle do not impact anxiety-like behavior in the open field (A) and light dark test (B-C) as well as home cage feeding following the NSF task (D-F)

## Abbreviations

DG =: Dentate Gyrus
FLX =: Fluoxetine
OVX =: Ovariectomized
EPM =: Elevated Plus Maze
NSF =: Novelty Suppressed Feeding
FST =: Forced Swim Test
Light Dark =: LD
Open Field =: OF
SSRI =: Selective Serotonin Reuptake Inhibitor
DCX =: Doublecortin

